# Chemical synthesis and purification of a non-hydrolyzable CTP analog CTPγS

**DOI:** 10.1101/2021.01.13.426546

**Authors:** M Rejzek, Tung B. K. Le

## Abstract

Slowly- or non-hydrolyzable analogs of ATP and GTP, for example adenosine 5’-(γ-thio)triphosphate (ATPγS) and guanosine 5’-(γ-thio)triphosphate (GTPγS), have been widely employed to probe the roles of ATP and GTP in biosystems, and these compounds are readily available from commercial sources. In contrast, cytosine 5’-(γ-thio)triphosphate (CTPγS) is not widely available commercially. The recent discovery of ParB as the founding member of a new class of CTPase enzyme (Osorio-Valeriano *et al*., 2019; Soh *et al*., 2019) and the possibility of multiple other undiscovered CTPases necessitate the use of the CTPγS to elucidate the roles of CTP hydrolysis in these systems. Here, we adapt a procedure for the synthesis of modified phosphoanhydrides (Hofer *et al*., 2015) to chemically synthesize and purify a milligram-scale of CTPγS.

## Introduction

A classical approach to the chemical synthesis of γ-thiophosphate analogs of nucleoside triphosphates is based on *S*-2-carbamoylethyl thiophosphate activation by diphenyl phosphorochloridate (Goody and Eckstein, 1971). The activated reagent was used to phosphorylate nucleoside diphosphates. The 2-carbamoylethyl protecting group was then removed by hydrolysis with sodium hydroxide and the product was purified by DEAE chromatography. This method has not yet been applied for preparation of CTPγS but the isolated yield for ATPγS was reported to be 25%.

A nucleoside diphosphate kinase mediated transfer of a thiophosphate group from a nucleoside 5′-(γ-thio)triphosphate to a nucleoside 5′-diphosphate was also reported (Goody et al., 1972). This method was applied to enzymatically synthesize CTPγS from ATPγS and CDP (Müller et al., 2000). A partially purified material generated by this transformation was treated with alkaline phosphatase to degrade all material devoid of terminal phosphorothioate, and with dithiothreitol to reduce any dimeric (CTPγS)_2_. Multistep HPLC purification yielded a low μmol quantity (about 10% overall yield) of pure CTPγS, which was characterized by UV and ^31^P NMR.

More recently, a highly versatile synthesis of nucleoside oligophosphates was reported (Hofer et al., 2015) based on protected phosphoramidites coupling to give mixed P^III^-P^V^ anhydrides. In the case of phosphorothioate analogs of nucleotides, the mixed P^III^-P^V^ anhydride was then oxidized by sulfur and the protecting group was removed.

Here, we adapt the Hofer et al., (2015) procedure to synthesize and purify a milligram-scale of CTPγS.

## Results and discussion

### Synthesis and purification of CTPγS

Briefly, the synthesis of CTPγS (**Scheme 1**) started from cytidine 5’-diphosphate (CDP) that was first converted into the corresponding tetrabutylammonium (TBA) salt. Care should be taken not to add excess tetrabutylammonium hydroxide. Ideally, CDP should be provided in a monoacidic form (2 x TBA). The exact number of TBA counterion was determined by ^1^H NMR to be cytidine 5’-diphosphate x 1.44 TBA salt. This salt was lipophilic enough to dissolve in DMF. The coupling was achieved with bis(9*H*-fluoren-9-ylmethyl)-diisopropylamidophosphite ((iPr)_2_NP(OFm)_2_) in the presence of 5-(ethylthio)-1*H*-tetrazole as an activator. The mixture was stirred for 20 minutes at room temperature and after that sulfur was added to oxidise the mixed P^III^-P^V^ anhydride. The bis-fluorenylmethyl-protected intermediate was then obtained by precipitation. Piperidine was used to remove the protecting groups and the crude product was precipitated again as a piperidinium salt. The crude product was purified by strong anion exchange chromatography (SAX) on a Poros 50 HQ using a gradient of ammonium bicarbonate to give CTPγS as an ammonium salt. The volatile buffer was removed by freeze drying. The purity of the final product was determined by SAX to be about 88%. The principal impurity was CDP formed by partial decomposition of CTPγS during the extensive freeze-drying procedure.

### CTPγS promotes the engagement between N-terminal domains of *Caulobacter crescentus* ParB

Previously, CTPγS has also been shown to promote the self-dimerization of the N-terminal domain (N-engagement) of ParB proteins (Jalal *et al*., 2020; Osorio-Valeriano *et al*., 2019; Soh *et al*., 2019). Here, we employed site-specific crosslinking of a purified ParB^*Ccres*^ (D141C C297S) variant by a sulfhydryl-to-sulfhydryl crosslinker bismaleimidoethane (BMOE) to test if our purified CTPγS can also promote the N-engagement of ParB. BMOE can covalently link symmetry-related D141C residues together if their inter-distance is within 8 Å. Crosslinking products in the presence or absence of nucleotides were analyzed by SDS-PAGE (**Figure 1**). We observed that CTP and CTPγS, but no other nucleotide triphosphate, enhanced the crosslinking of ParB^*Ccres*^ (D141C C297S) (**Figure 1**), consistent with the previous findings that CTP hydrolysis is not required for N-engagement (Jalal *et al*., 2020; Soh *et al*., 2019).

**Figure 1.**
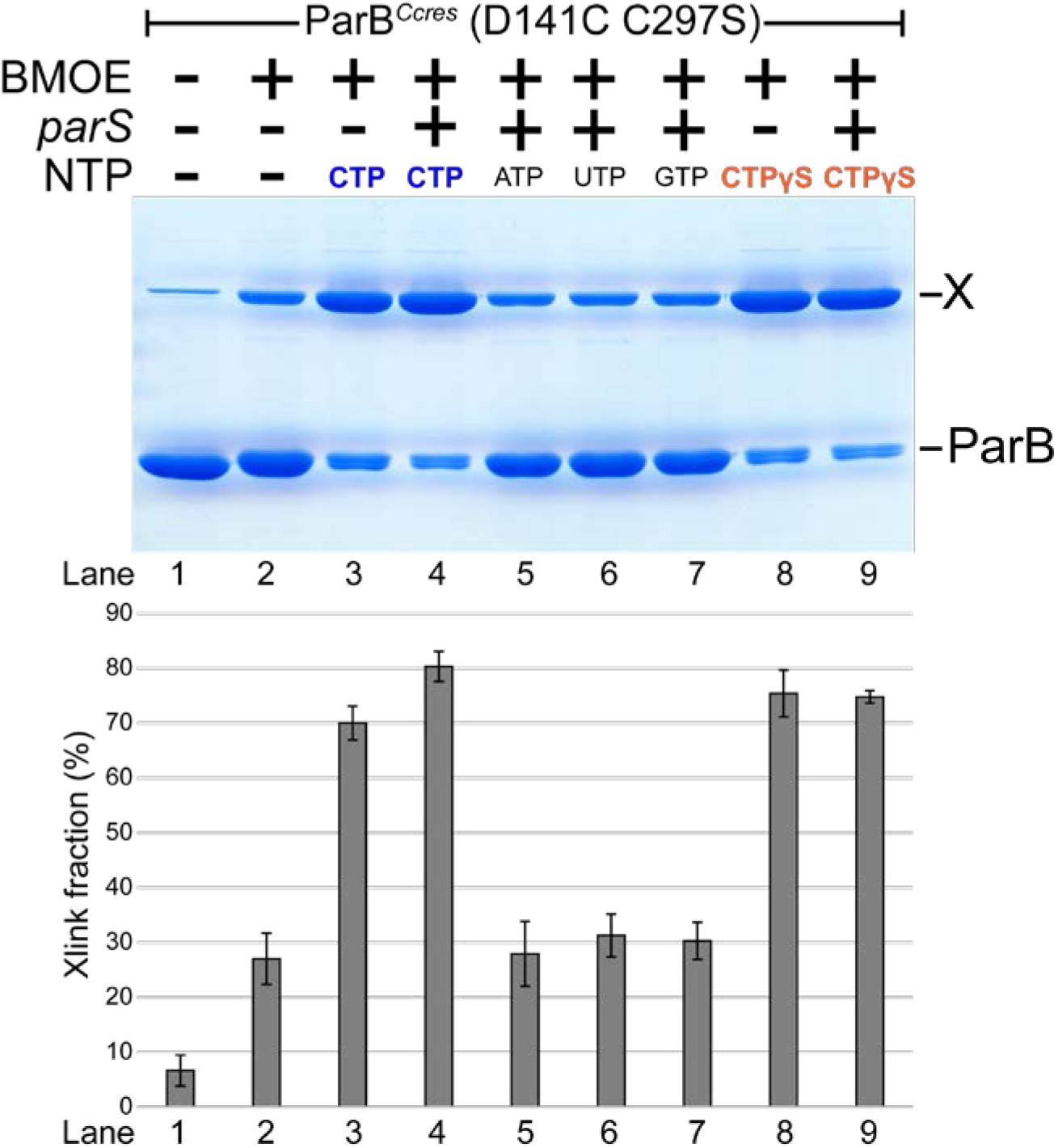
SDS-PAGE analysis of BMOE crosslinking products of *C. crescentus* ParB (D141C C297S) in the presence of CTPγS or other nucleotides triphosphates. A 50 µL reaction that contained 8 µL of ParB (D141C C297S) ± 1 mM NTP ± 0.5 µM 22-bp *parS* DNA in a binding buffer [10 mM Tris-HCl pH 7.4, 100 mM NaCl, 1 mM MgCl_2_] was incubated for 5 minutes at 22°C. BMOE (1 mM final concentration) was added and the mixture was quickly mixed by vortexing. SDS-PAGE sample buffer was then added to quench the crosslinking reaction. Samples were heated to 50°C and analyzed by SDS-PAGE. Crosslinked fraction was quantified from triplicated experiments ± standard deviation.

## Conclusions

Here, we report a straightforward procedure for the chemical synthesis and purification of a milligram-scale of a non-hydrolyzable CTP analog (CTPγS). Improved availability of CTPγS will facilitate research by the community into ParB/ParB-like proteins and other CTPases enzymes.

### Experimental procedures Materials and equipment

Sulfur (Cat# 213292-10G) was purchased from Sigma Aldrich, UK. 5-(Ethylthio)-1*H*-tetrazole (Cat# 493805-2G) was purchased from Sigma Aldrich. Cytidine 5’-diphosphate (Cat# C9755-100MG) was purchased from Sigma Aldrich as a sodium salt and converted into its tetrabutylammonium (TBA) salt by ion exchange on Dowex 50WX8 (H^+^), followed by neutralization with TBA hydroxide (Cat# 178780-50ML) and subsequent lyophilization. Bis(9*H*-fluoren-9-ylmethyl)-diisopropylamidophosphite (Cat# SC-0057, (iPr)_2_NP(OFm)_2_) was purchased from SiChem, Germany. Bis-maleimidoethane (BMOE) powder was purchased from ThermoFisher (Cat# 22323) and was dissolved in DMSO to 20 mM. NMR spectra were recorded on a Bruker Advance III 400 MHz spectrometer. Chemical shifts of ^1^H NMR signals were recorded in D_2_O and were reported with respect to residual solvent peak at δ_H_ 4.79 ppm. 31P{1H} NMR spectra were recorded with ^1^H‐decoupling on Bruker 162 MHz spectrometer in D_2_O. High resolution accurate mass spectra were obtained using a Synapt G2-Si Q-Tof mass spectrometer using negative electrospray ionisation. HPLC purification was performed on a Dionex Ultimate 3000 instrument equipped with an UV/vis detector. Lyophilization was performed with a Labconco FreeZone Benchtop Freeze Dryer with PTFE Coil. An Eppendorf 5810R benchtop centrifuge was used for centrifugation.

### Synthesis of cytosine 5’-(γ-thio)triphosphate (CTPγS)

Cytidine 5’-diphosphate (CDP) (8.34 mg, 20.7 μmol) in ultrapure Milli-Q water (1 mL) was applied on a column containing Dowex 50WX8 (H^+^) (1 mL, pre-washed with HCl and Milli-Q water to neutral pH). The column was eluted with Milli-Q water (10 mL) while monitoring the pH, and elution was stopped when the pH returned to neutral. Tetrabutylammonium hydroxide solution (∼40% weight in H_2_O, 41 μL) was further diluted with water to a final volume of 1 mL. Out of 1 mL, 300 μL was added to the eluted CDP while monitoring the pH, the final pH was ∼7.0. Care was taken not to use an excess of TBA hydroxide. The sample was subsequently freeze-dried. The accurate number of counterions was determined by ^1^H NMR (D_2_O) to be cytidine 5’-diphosphate x 1.44 nBuN^+^ salt (**Figure S1**).

^1^H NMR (400 MHz, D_2_O) δ 8.11 (d, J = 7.9 Hz, 1H), 6.20 (d, J = 7.8 Hz, 1H), 5.90 (d, J = 3.7 Hz, 1H), 4.33-4.28 (m, 2H), 4.26-4.21 (m, 2H), 4.16-4.11 (m, 1H), 3.17-3.11 (m, 11.5H), 1.63-1.55 (m, 11.5H), 1.34-1.25 (m, 11.6H), 0.89 (t, J = 7.4 Hz, 17.4H).

Next, cytidine 5’-diphosphate x 1.44 nBu_4_N^+^ (10.67 mg, 14.2 μmol) was dissolved in 0.5 mL of dimethylformamide (DMF). To this solution, (iPr)_2_NP(OFm)_2_ (13.6 mg, 26.2 μmol) was added and immediately after its dissolution, 5-(ethylthio)-1*H*-tetrazole (4.8 mg, 36.8 μmol) was added. The mixture was stirred for 20 minutes at room temperature. After that sulfur (3.0 mg, 94.7 μmol) was added and the mixture was mixed vigorously by vortexing for 2 minutes. The bis-Fm-protected intermediate was then obtained by precipitation with Et_2_O/hexane 5:1 (7.5 mL) and hexane (2 mL), and pelleted by centrifugation (184 x g, at 4°C for 5 minutes). The resulting pellet was washed with Et_2_O and dried under a stream of nitrogen gas and briefly in vacuum. The precipitate was then dissolved in 0.5 mL of DMF. Piperidine (25 μL, 21.55 mg, 253.1 μmol, 5% v/v) was added and the mixture was stirred for 15 minutes. The final product was precipitated by adding 3.5 mL of 1:5 DCM/Et_2_O 1:5. The mixture was centrifuged at 184 x g for 5 minutes at 4°C. The pellet was washed with Et_2_O and dried under a nitrogen stream to give crude CTPγS as a piperidinium salt. The sample was stored at −80°C, which is stable for approximately 2 months, until the purification step.

^31^P{^1^H} NMR (162 MHz, D_2_O) δ 34.2 (d, J = 29.5 Hz), −11.4 (d, J = 19.7 Hz), −23.6 (dd, J = 29.5, 19.7 Hz). The ^31^P NMR spectrum (**Figure S2**) is in good agreement with data published previously, in which CTPγS was synthesized enzymatically (Müller *et al*., 2000).

### Purification of CTPγS

Next, the sample was dissolved in 1 mL of Milli-Q water and filtered through a 0.45 μm PTFE disk filter. CTPγS was purified by strong anion exchange chromatography (SAX) on a Poros 50 HQ column (10 x 100 mm) at a flow rate of 7 mL/min and UV detection at 265 nm. The elution solvents were 5 mM ammonium bicarbonate (solvent A) and 250 mM ammonium bicarbonate (solvent B). The elution started with 0% solvent B over 2.5 minutes, 100% solvent B over 7.5 minutes, hold for 4.5 minutes, 0% solvent B over 1 minute, and finally the column was equilibrated at 0% solvent B for 1 minute. The target compound (CTPγS) was eluted as the last peak at *R*_t_ = 12.3 min. Combined CTPγS-containing fractions were freeze-dried to give the final product as an ammonium salt (2.15 mg, yield: ∼26.7%, final purity by HPLC: ∼88%) (**Figure S3**).

^1^H NMR (400 MHz, D_2_O) δ 8.17 (d, J = 7.8 Hz, 1H), 6.31 (d, J = 7.8 Hz, 1H), 5.99 (d, J = 4.2 Hz, 1H), 4.44 (dd, J = 4.9, 4.9 Hz, 1H), 4.40-4.38 (m, 1H), 4.37-4.26 (m, 3H) (**Figure S4**).

^31^P{^1^H} NMR (162 MHz, D_2_O) δ 38.7 (bd, J = 26.7 Hz), −11.4 (d, J = 20.3 Hz), −23.5 to −24.0 (m) (**Figure S5**).

HR-MS (ESI^-^) for C_9_H_15_N_3_O_13_P_3_S^-^ m/z calcd.: 497.9544 found: 497.9544 [M-H]^-^ (**Figure S6**).

### Construction of pET21b::*Ccres parB (D141C C297S)-his*_*6*_

The DNA fragment containing *parB (D141C C297S)* was chemically synthesized (gBlocks, IDT). The NdeI-HindIII-cut pET21b plasmid backbone and *parB (D141C C297S)* gBlocks fragments were assembled together using a 2x Gibson master mix (NEB). Gibson assembly was possible owing to a 23-bp sequence shared between the NdeI-HindIII-cut pET21b backbone and the gBlocks fragment. The resulting plasmids were sequenced verified by Sanger sequencing (Genewiz, UK).

### Purification of *Ccres* ParB (D141C C297S)-His_6_

The same procedure as previously described (Jalal et al., 2020) was used to purify the ParB (D141C C297S) variant. Purified protein was buffer-exchanged and stored in a storage buffer supplemented with TCEP [100 mM Tris-HCl pH 7.4, 300 mM NaCl, 10% (v/v) glycerol, and 1 mM TCEP].

### *In vitro* crosslinking assay using a sulfhydryl-to-sulfhydryl crosslinker bismaleimidoethane (BMOE)

A 50 µL mixture of 8 µM ParB (D141C C297S) ± 1 mM NTP ± 0.5 µM 22-bp *parS* DNA was assembled in a reaction buffer [10 mM Tris-HCl pH 7.4, 100 mM NaCl, and 1 mM MgCl_2_] and incubated for 5 minutes at room temperature. BMOE (1 mM final concentration from a 20 mM stock solution) was then added, and the reaction was quickly mixed by three pulses of vortexing. SDS-PAGE sample buffer containing 23 mM β-mercaptoethanol was then added immediately to quench the crosslinking reaction. Samples were heated to 50°C for 5 minutes before being loaded on 12% Novex WedgeWell Tris-Glycine gels (ThermoFisher). Protein bands were stained with an InstantBlue Coomassie solution (Abcam) and band intensity was quantified using Image Studio Lite version 5.2 (LI-COR Biosciences). The crosslinked fractions were averaged, and their standard deviations from triplicated experiments were calculated in Excel.

## Acknowledgments

This work was funded by BBSRC grant (BBS/E/J/000PR9791) to T.B.K.L.

---

**Scheme 1.**
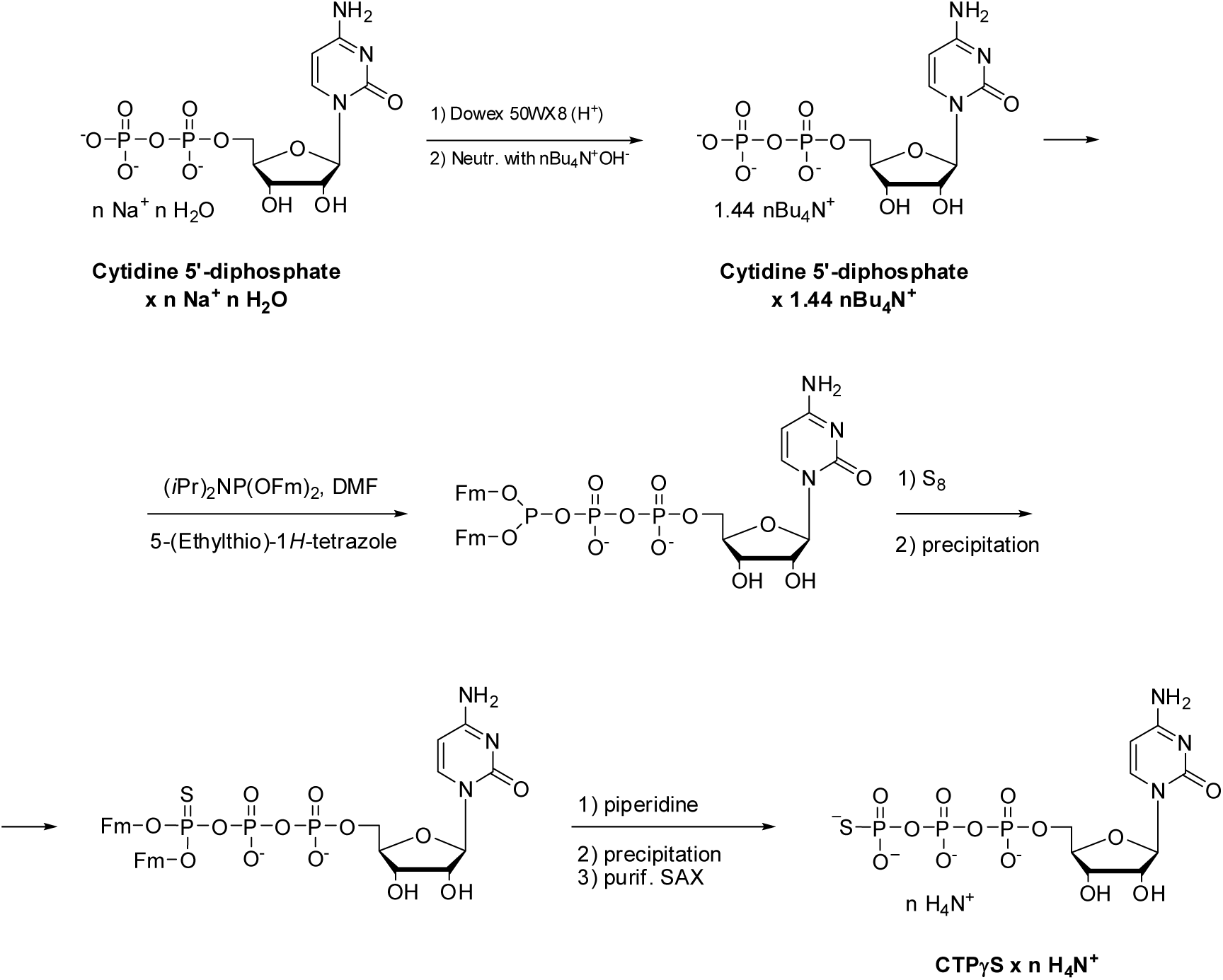
Synthesis of cytosine 5’-(γ-thio)triphosphate (CTPγS)

## Supporting Information

**Figure S1.**
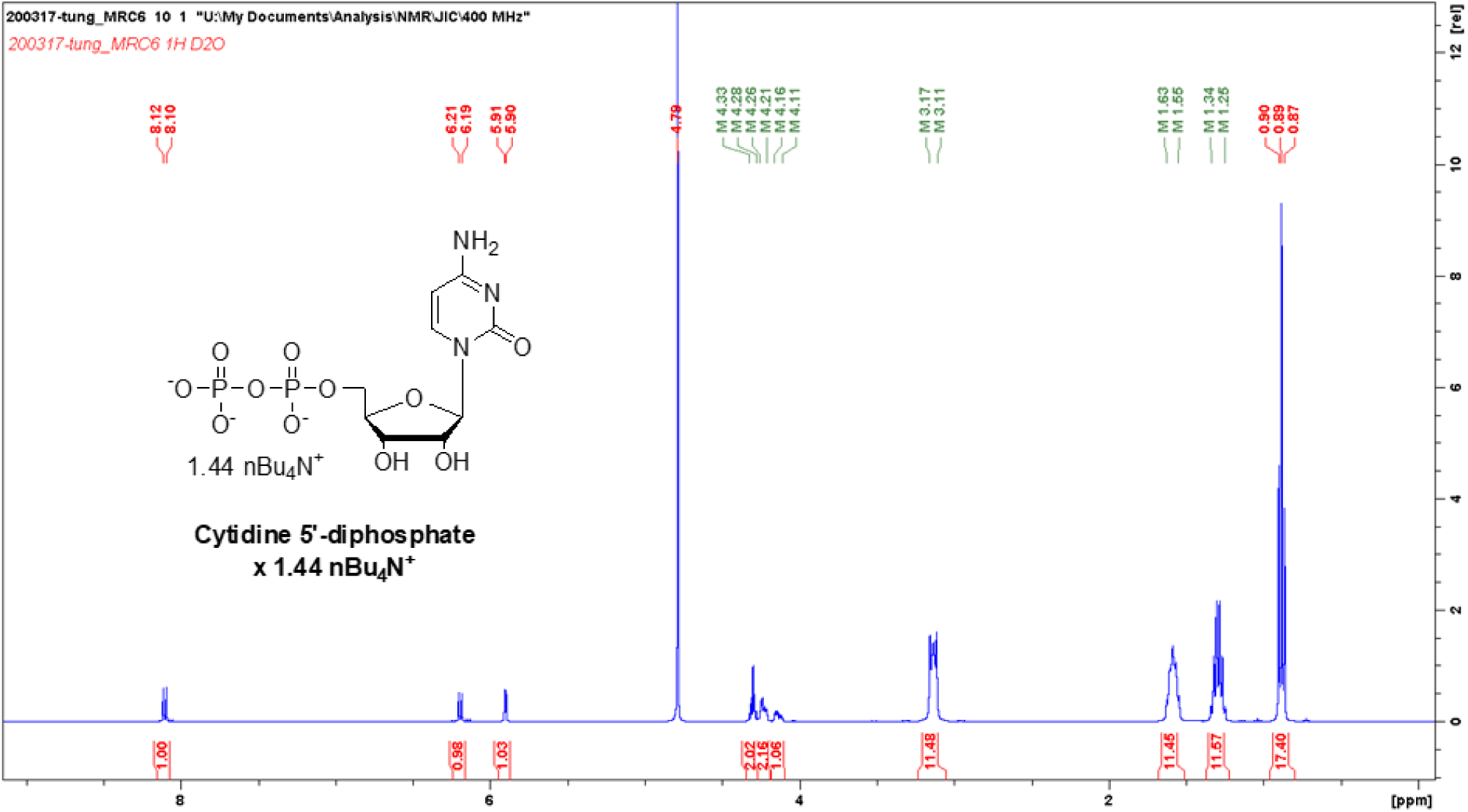
Cytidine 5’-diphosphate x 1.44 nBuN^+^salt:^1^H NMR (400 MHz, D_2_O)

**Figure S2.**
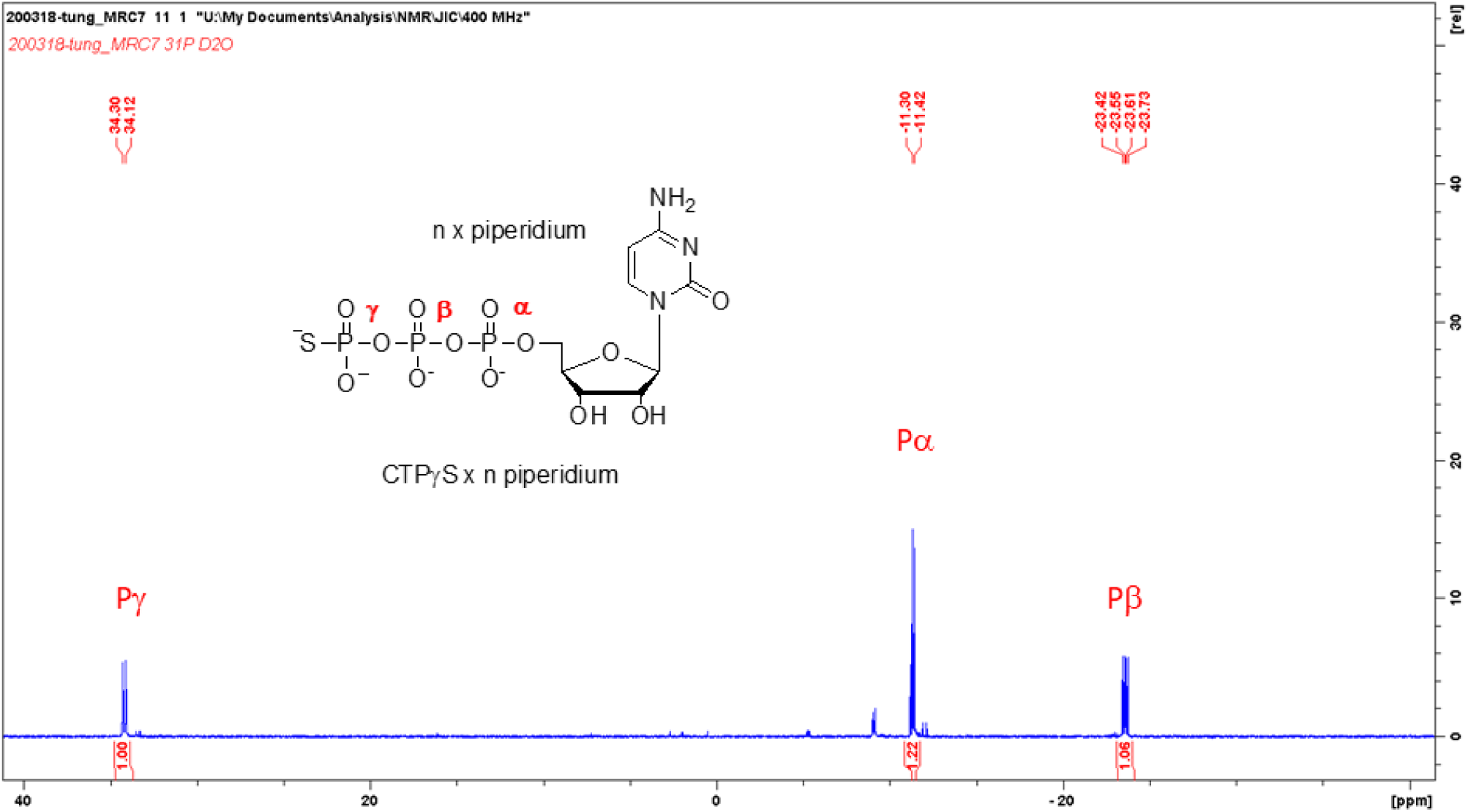
CTPγS x n piperidium salt: ^31^P{1H} NMR (162 MHz, D_2_O)

**Figure S3.**
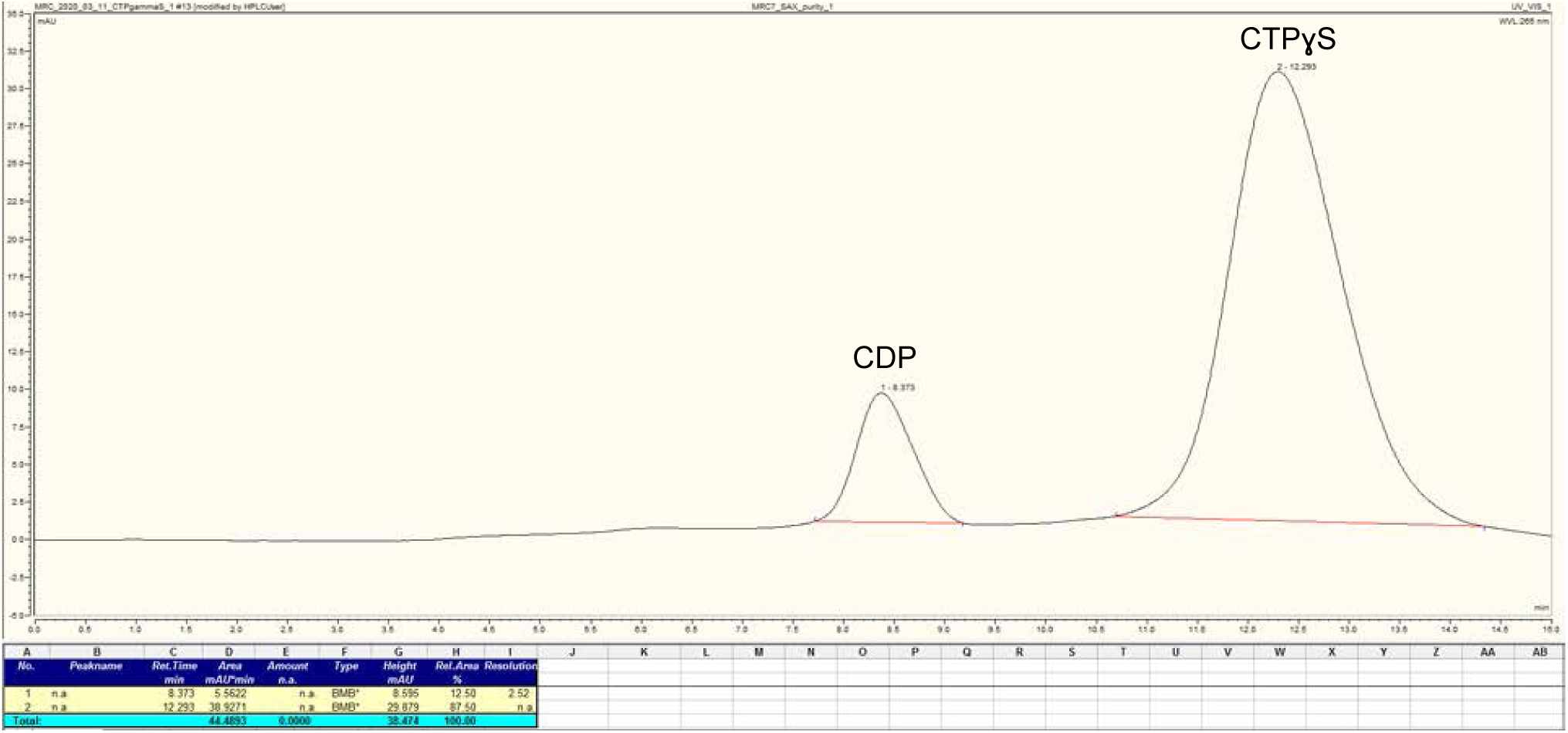
HPLC (Poros 50 HQ, ammonium bicarbonate, 5 to 250 mM) Retention time: 12.3 min. Final purity (HPLC): 88%.

**Figure S4.**
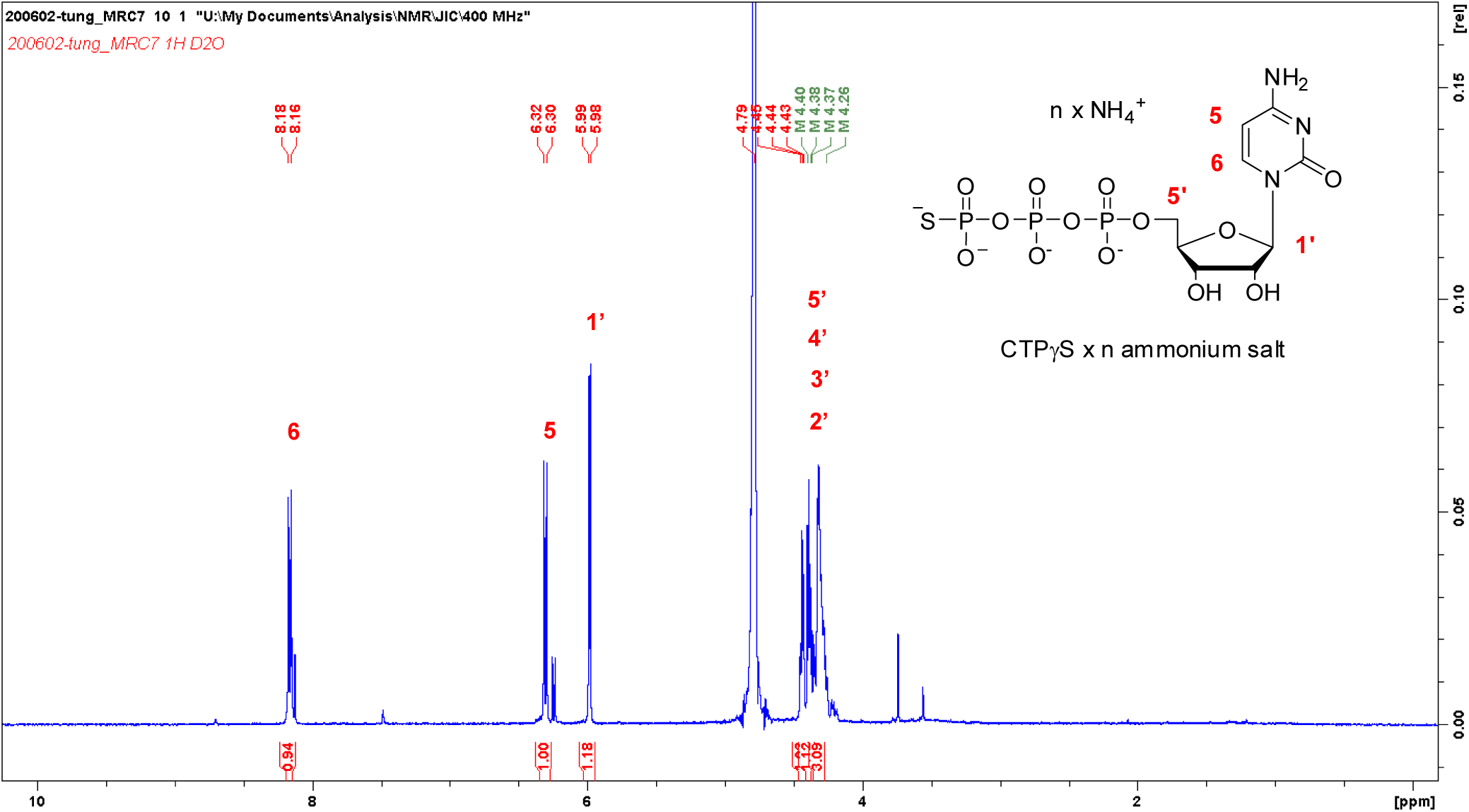
CTPγS x n ammonium salt: ^1^H NMR (400 MHz, D_2_O)

**Figure S5.**
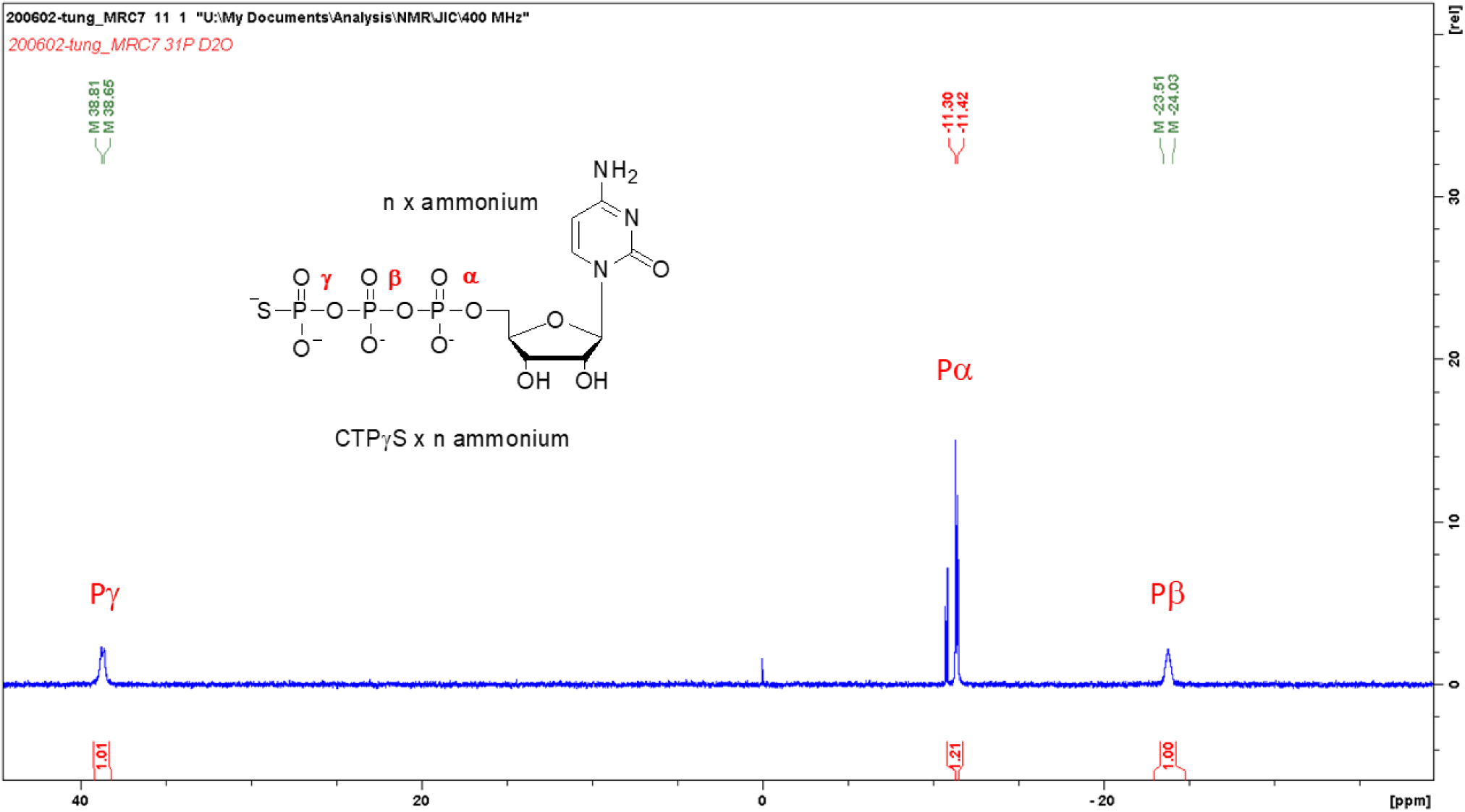
CTPγS x n ammonium salt: ^31^P{1H} NMR (162 MHz, D_2_O)

**Figure S6.**
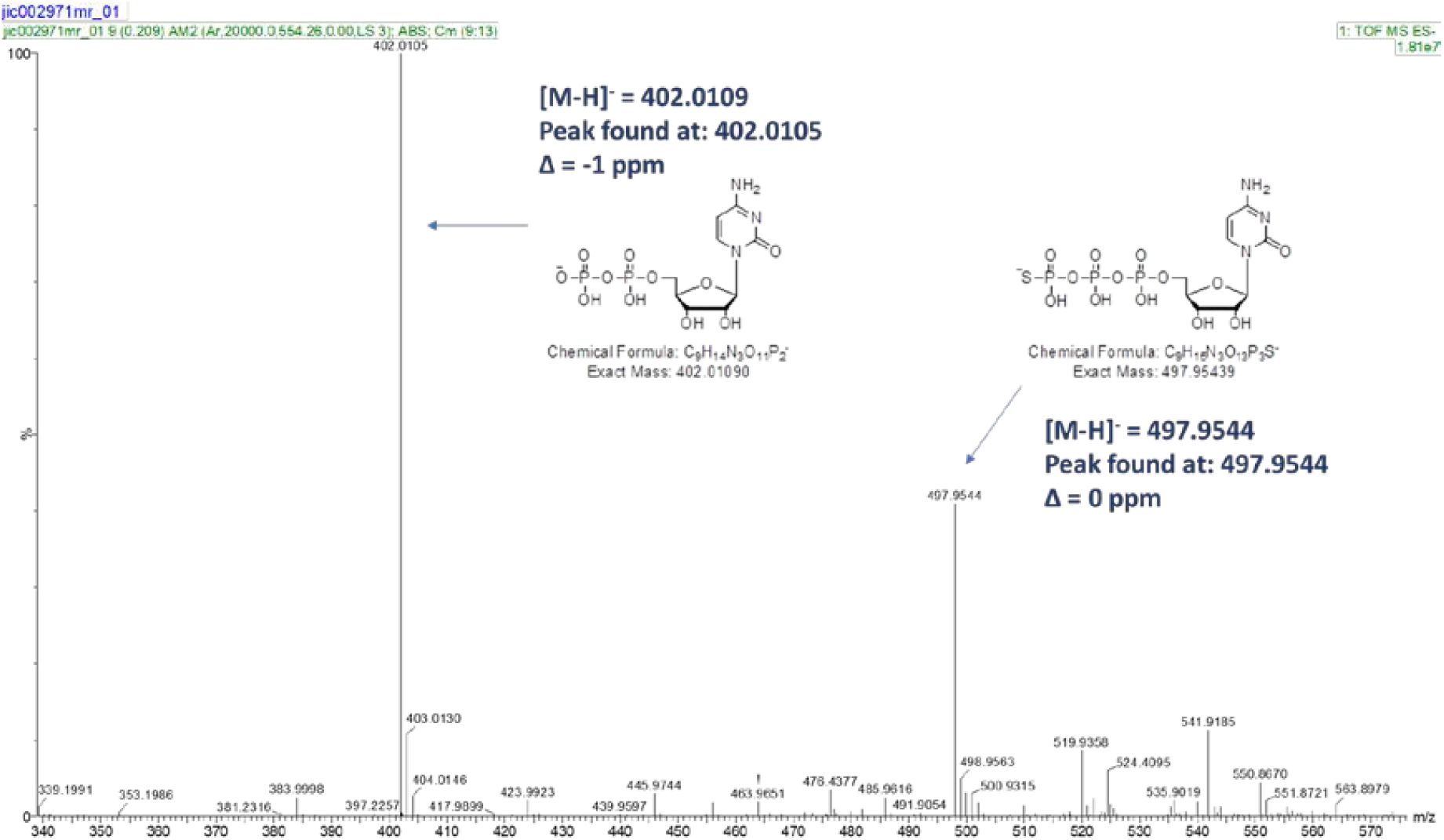
CTPγS: HR-MS (ESI^-^)

